# Inferring Effective Networks of Spiking Neurons Using a Continuous-Time Estimator of Transfer Entropy

**DOI:** 10.1101/2024.09.22.614302

**Authors:** David P. Shorten, Viola Priesemann, Michael Wibral, Joseph T. Lizier

**Author notes:** ^u^.

## Abstract

When analysing high-dimensional time-series datasets, the inference of effective networks has proven to be a valuable modelling technique. This technique produces networks where each target node is associated with a set of source nodes that are capable of providing explanatory power for its dynamics. Multivariate Transfer Entropy (TE) has proven to be a popular and effective tool for inferring these networks. Recently, a continuous-time estimator of TE for event-based data such as spike trains has been developed which, in more efficiently representing event data in terms of inter-event intervals, is significantly more capable of measuring multivariate interactions. The new estimator thus presents an opportunity to more effectively use TE for the inference of effective networks from spike trains, and we demonstrate in this paper for the first time its efficacy at this task. Using data generated from models of spiking neurons — for which the ground-truth connectivity is known — we demonstrate the accuracy of this approach in various dynamical regimes. We further show that it exhibits far superior inference performance to a pairwise TE-based approach as well as a recently-proposed convolutional neural network approach. Moreover, comparison with Generalised Linear Models (GLMs), which are commonly applied to spike-train data, showed clear benefits, particularly in cases of high synchrony. Finally, we demonstrate its utility in revealing the patterns by which effective connections develop from recordings of developing neural cell cultures.

## I. INTRODUCTION

For many of the complex systems that scientists are most interested in, our ability to record high-fidelity data from the numerous components of these systems is improving rapidly. For instance, the number of biological neurons that can be simultaneously recorded from is increasing exponentially, with a doubling rate of around six to seven years [1, 2], while spatial resolution is increasing at a similar rate [3]. The process of drawing scientific insight from this flood of data is, however, not always straightforward [4].

The inference of effective network models [5] from high-dimensional time-series data has become a popular and productive technique for reducing the complexity of this class of data. Such datasets often consist of millions of (or far more) individual data points [6]. The inference of effective networks aims to produce a minimal system model from the data, by finding the smallest set of system source components capable of explaining the activity of each target component [7]. As such, it compresses the large number of data points down to a single directed network diagram describing the relationships between components of the system, thus facilitating the interrogation of the data at hand.

In this work, we specifically focus on the inference of effective networks for time-series data of spiking neural activity, summarised by the timestamps of action potentials (spikes) at each node in the network. Much research has addressed network relationships using Transfer Entropy (TE)[8, 9], a widely accepted measure of information transfer, suitable for mapping information flows in neural systems [10]. This has incorporated exploring [11–14] and applying TE-based methods to *in vitro* [15–21] and *in vivo* [22] recordings of spiking activity. All of this work has estimated TE in a pairwise fashion, producing what are often referred to as *directed functional* (as opposed to *effective*) networks [5].

The main contribution of this work is to marry a recently-proposed continuous-time estimator of TE for spike trains [23] with an existing greedy effective network inference algorithm. The greedy approach iteratively adds sources to a parent set for each target in a greedy fashion, choosing the source with the highest conditional TE at each step until no candidate sources with (statistically significant) non-zero TE remain [7, 24–27].

The main obstacle that has prevented the inference of effective networks from spike trains using TE has been the manner in which the traditional method of estimating TE on spike train data – which uses long sequences of time bins to represent temporal spiking patterns – causes a rapid increase in the dimensionality as we add conditioning processes [23].

The recent development of a continuous-time estimator of TE for event-based data [23] bypasses the dimensionality problems of the discrete-time estimator. Specifically, as it uses inter-spike intervals to efficiently represent the temporal spiking patterns, it is capable of capturing dependencies over relatively long periods of time (on the order of seconds [28]), with no loss of time precision. This makes the estimation of TE with sizeable conditioning sets feasible, supporting efficient inference of effective networks.

In this paper, we bring the greedy network inference algorithm and the continuous-time TE estimator together for the first time. We validate the efficacy of this combination on synthetic examples, where the underlying causal network is recovered by the model. We further compare its efficacy against generalised linear models [29] and a convolutional neural network based approach [30], finding its performance to be highly competitive. We finally demonstrate its ability to uncover biological insight by inferring the effective networks of developing cell cultures of dissociated cortical rat neurons [31].

## II. RESULTS

In this section we apply the greedy TE-based effective network inference algorithm [7] in conjunction with the continuous-time TE estimator for event-based data [23]. Please see Sec. IV A for details of the operation of the greedy algorithm, along with a description of a few minor changes that were made for the application to event-based data. Sec. IV D summarises the TE estimation approach used.

The first three subsections of this section focus on the inference of simulated spiking networks for which the ground truth connectivity is known. We must emphasize, however, that in general we do not expect the effective networks inferred by TE to align with the causal structure [7]. Whilst the effective networks always provide a useful model for interpreting the directed relationships in the system, it is only under certain specific conditions that we expect them to match the causal structure, most importantly full observability of the nodes involved in the dynamics, and under certain assumptions such as faithfulness and the causal Markov property [32, 33]. Therefore, we evaluate the performance of the network inference scheme by comparing the inferred network to this ground truth under these idealised conditions, since this provides an important validation of the output of the inference when this match can be expected. In order to measure the accuracy of the inference scheme, we make use of the commonly-employed classification metrics of *recall* and *precision*. They are defined as:

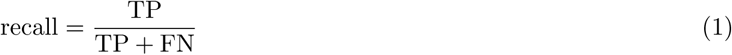

and

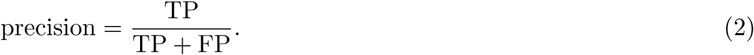

Here, TP is the number of true positives, FP is the number of false positives and FN is the number of false negatives. In the context of network inference, recall can be interpreted as the proportion of true connections that were predicted by the algorithm. Precision, on the other hand, is the proportion of predicted edges that are true edges.

The final subsection of the results focuses on the application of the estimator to the spike times from recordings of cultures of dissociated rat cortical neurons. This provides a demonstration of the utility of this approach for extracting insights from biological data.

### A. Inference at varying levels of synchrony

We constructed networks of Leaky-Integrate-and-Fire (LIF) [34] neurons with alpha synapses [35]. These networks were composed of 30 excitatory and 20 inhibitory neurons. Each neuron had exactly three excitatory and two inhibitory sources, where these sources were selected randomly from the respective sets. By varying the ratio of inhibitory to excitatory connection strength *g*, we could vary the level of synchrony within the networks. We ran simulations for three different levels of synchrony, which we refer to as “low” (*g* = 3), “medium” (*g* = 1.5) and “high” (*g* = 1). Varying the relative strength of inhibitory connections across the excitation-inhibition balance threshold is a known method for adjusting the degree of synchrony in these networks [36]. Please see Sec. IV F for full details on these network models. It is also worth noting that the level of synchrony present in these networks (even at “high” synchrony) is far lower than in the biological data examined in Sec. II D.

The combination of the greedy inference algorithm and the continuous-time estimator was applied to these networks, and the resulting inferred networks were compared against the ground truth for varying numbers of target spikes available to the estimator: 100, 300, 500, 1000, and 3000 (extra runs at 5000 target spikes were included for the high synchrony network, as the recall rose more slowly in this case). 10 independent simulations of the network model were run for each level of synchrony and the algorithm was applied to each run for each number of target spikes, although it was only applied to the first 5 simulations at 3000 spikes and the first 3 simulations at 5000 spikes due to the high computational requirements.

The precision and recall of the resulting inferences was calculated and is plotted in Fig. 1 for the different dynamical regimes and numbers of target spikes.

**FIG. 1:**
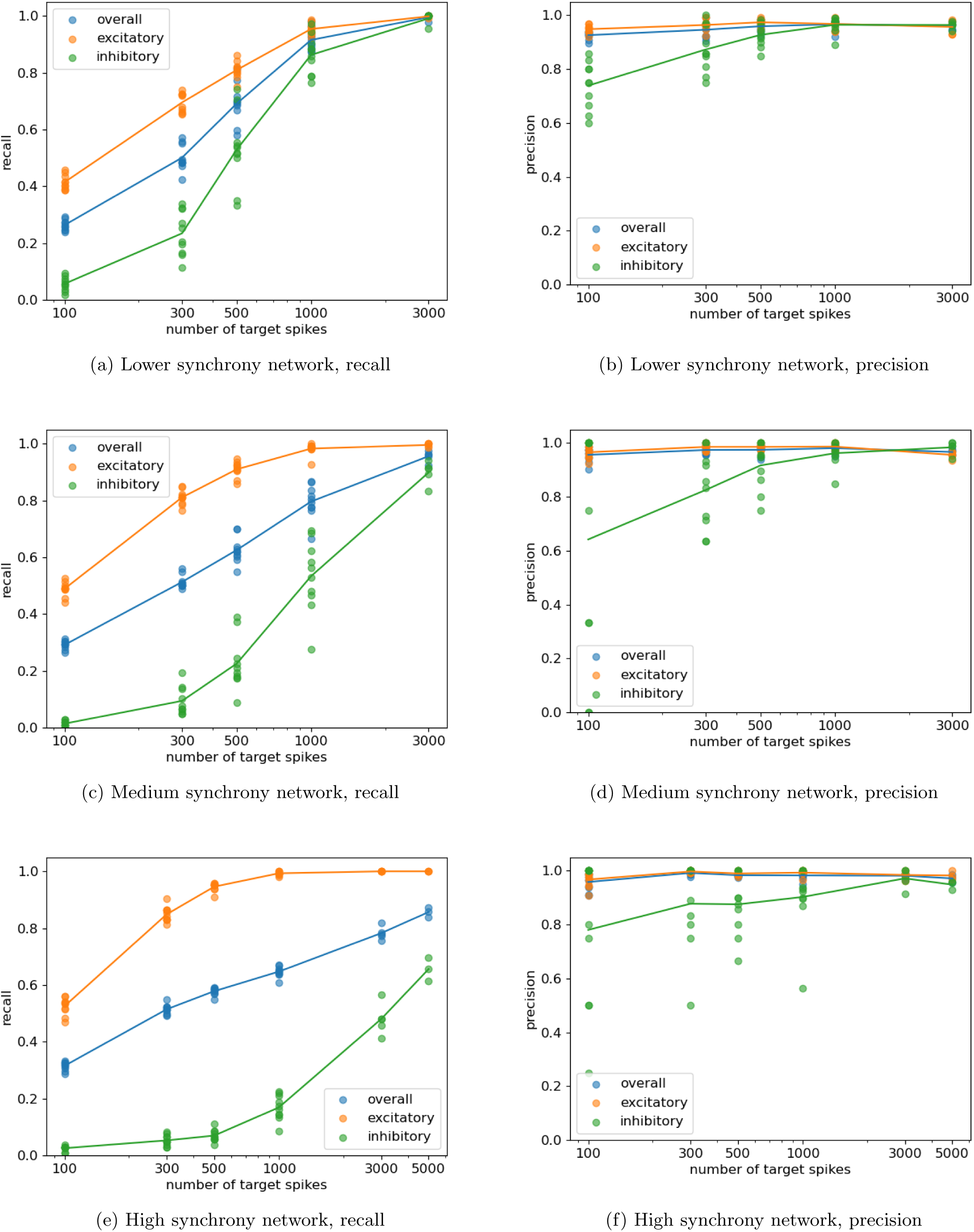
Plots showing the resulting precision and recall from running the network-inference scheme on networks of LIF neurons composed of 30 excitatory neurons and 20 inhibitory neurons. Each neuron has exactly 3 excitatory and 2 inhibitory inputs, chosen randomly. The ratio of the inhibitory to excitatory connection strength was varied in order to change the level of synchrony in the network. Plots are shown for three different synchrony levels. Each plot contains points for each experiment the precision and recall for the inhibitory and excitatory sources separately as well as for their overall weighted average. The lines pass through the means of these points.

In the results shown in Fig. 1, we see that the algorithm exhibits high precision for all combinations of dynamical regime and number of target spikes. The precision only drops below 0.9 where the recall is very low, i.e. when few links are inferred. This demonstrates the high confidence with which it predicts links — a very low proportion of the predicted links turn out to be false positives – which can also be seen as a conservative approach.

In these plots, the recall begins low, but rises rapidly with the increase in the number of target spikes available. In the case of low network synchrony (Fig. 1a), we observe that, by 3000 target spikes, the recall has risen to nearly one. As the precision is also nearly one at this number of target spikes, the networks are being inferred nearly perfectly. Taken in conjunction with the apparent trends towards converging on perfect inference as the number of spikes increases for the other regimes, this suggests evidence for validation that the approach provides a consistent inference of the underlying network under the aforementioned idealised conditions.

By comparing Figs. 1a, 1c and 1e, we can observe that the achieved overall recall drops as the level of synchrony is increased. This is entirely driven by a drop in the recall on inhibitory connections. In fact, we observe a small increase in the recall on excitatory connections. This drop in recall is due to the increased complexity in the nature of the statistical relationship between the activity of a given target and an inhibitory source in the case of high synchrony. When the populations are highly synchronous, all of the cells spike close together, so the firing of an inhibitory source can become positively correlated with the firing of its target, when considering the purely pairwise relationship between the source and target. Note that the genuine relationship that we are seeking to infer here is the negative correlation in firing resulting from inhibition. It is only when conditioned on the target’s excitatory sources (which becomes possible with more target spikes observed), for any given firing pattern of the excitatory sources, the firing of the inhibitory source is associated with a decrease in the probability of the firing of the target for that given pattern, allowing the inhibitory source to be identified. Crucially the precision remains high despite the spurious pairwise correlations that appear in the highly synchronous regime, due to the conditioning in the multivariate approach removing such redundant sources being included in the inference. Fig. 3f shows an ROC curve for the use of a purely pairwise approach on this same high-synchrony example for 1000 target spikes, which will be discussed in Sec. II C. We see that it cannot achieve a high true positive rate without a substantial increase in the false positive rate (and thus a decrease in the precision). This highlights the necessity of using a full multivariate approach when dealing with highly-synchronous neural populations.

In order to compare the performance of the proposed scheme with an existing network inference approach, we ran the recently-proposed CoNNECT [30] algorithm on the spike times from the simulations. This approach makes use of pre-trained convolutional neural networks and has been demonstrated to be competitive when compared with other existing network inference algorithms for spike trains. In order to perform this inference, we made use of the associated web-app provided by the authors [37]. The resulting precision and recall plots are shown in Fig. 8 of Appendix A. We consistently see that, for any given combination of number of target spikes and dynamical regime, the proposed approach is able to achieve both higher precision and higher recall. In particular, the precision of the results of the CoNNECT inference is particularly low, being largely in the region 0.1-0.2.

We also compared the performance of the proposed approach with a Generalised Linear Model (GLM), which is a popular approach for modelling spiking neural data [29, 38, 39], including for inferring connectivity [40, 41]. We closely followed previous work [40] which demonstrated the use of these models for connectivity inference, with a few minor differences, as specified in Sec. IV H. The resulting precision and recall plots are shown in Fig. 9 of Appendix A. The GLM approach exhibits markedly lower precision than the proposed TE-based approach, except for precision on inhibitory sources for very low numbers of available spikes in the target spike trains, where our more conservative approach is inferring very few links. Moreover, in the case of high synchrony, the precision of the proposed approach is far superior, whereas the very low precision of the GLM approach results in almost half of all possible connections being inferred as present, rendering those inferred network models far less useful. For low numbers of target spikes, the GLM approach is able to achieve better recall. However, this is always at a cost of significantly lower precision, and any advantage in recall for the GLM approach disappears as the number of target spikes is increased above 1000 (except in the high-synchrony case). Moreover, these results were achieved by carefully tuning certain parameters of the GLM model setup in order to maximise performance on these specific examples. Importantly, this parameter tuning was done using the known ground-truth. In real-world inference applications we do not have access to such a ground truth. Using sub-optimal parameters can have a dramatic effect on the performance of this approach. Fig. 10 shows how the precision deteriorates even further when a larger bin size Δ*t* is used ((40 ms as oppose to 20 ms)). Fig. 11 shows a similar deterioration in performance when we more closely mirror the original implementation by Song et. al. [40] (i.e. by increasing the number of knots used in the spline basis functions to include all 16 knot locations in the original). Interestingly, the better recall that our approach shows for excitatory versus inhibitory sources is reversed in the GLM approach.

### B. Inference at varying levels of stimulus regularity

In Sec. II A neurons were always provided with an independent Poisson stimulus. However, we can vary the properties of this stimulus in order to mimic different plausible inference scenarios. For instance, simulations constructed with a fully regular stimulation of the neurons provide us with an example of dynamics with no hidden sources of variability. That is, it is possible to perfectly predict the dynamics of the neuron based on its past and the past of those neurons connecting to it. Moreover, the system is completely deterministic. As we move towards the semi-regular and fully random Poisson stimuli, we are modelling increasing amounts of hidden activity or noise within the system, which prevents perfect predictability of the future state of the units. Full determinism can make the inference of networks from time series more difficult [32]. Conversely, very large amounts of noise can make it challenging to detect a comparatively weak relationship between two nodes. As such, it is important that both potentially problematic ends of this spectrum are tested.

We constructed model networks as in Sec. II A, however, this was done at a single level of ratio of inhibitory to excitatory connection strength. The same ratio (*g* = 3) used to produce the low synchrony runs was used. Instead, we varied the nature of the stimulus provided to each neuron, providing them with a regular, semi-regular or Poisson (fully random) stimulus. The regular stimulus was composed of spike times placed at a fixed interval. There was slight variation in this interval between neurons, in order to prevent the network settling into a simple, fixed, pattern. The semi-regular stimulus was similar to the regular stimulus, but with the addition of a small amount of Gaussian noise. Note that a few other minor simulation parameters had to be changed from the simulations used in Sec. II A in order to ensure numerical stability. See Sec. IV F for a full specification of these differences.

Fig. 2 shows plots of precision and recall, for different numbers of observed spikes in the target, for these different levels of stimulus regularity. We observe that the recall increases slightly when moving from the regular to the semiregular stimulus, but then drops as we move to the Poisson stimulus. By contrast, the precision exhibited a slight increase with increasing irregularity. This is likely because the increasing irregularity reduces the correlations between true and false sources, thereby decreasing the likelihood of false positives. In all three cases, good performance is achieved at 3000 target spikes, with overall precision and recall both being around 0.9.

**FIG. 2:**
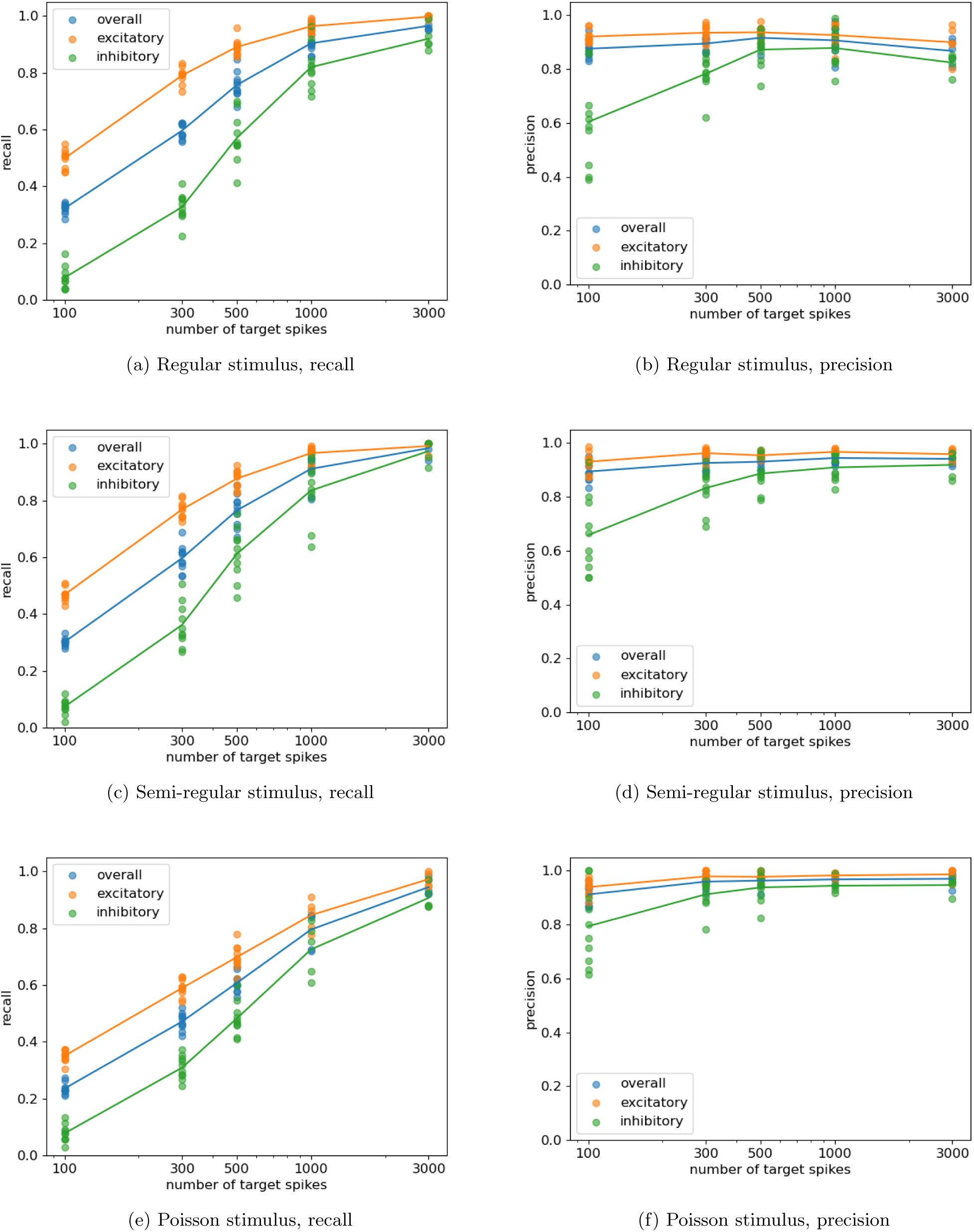
Plots showing the resulting precision and recall from running the network-inference scheme on networks of LIF neurons composed of 30 excitatory neurons and 20 inhibitory neurons. The regularity of the stimulus provided to each neuron was varied.

These results demonstrate that the proposed combination of estimator and network inference scheme is capable of successful inference at various levels of determinism or unobserved noise.

### C. Comparing the greedy algorithm to pairwise inference

Recent work [42] has highlighted the improvements that can be gained when performing network inference using TE in its full multivariate sense, via the greedy algorithm, as opposed to the simpler pairwise approach. The pairwise approach operates by only checking for a statistically significant non-zero TE value between each source-target pair, without taking into account the other processes in the system. It is generally found that the full multivariate approach tends to exhibit much higher precision, as, among other reasons, it is able to distinguish true sources (which provide information about the target even when conditioned on all other system components) from spurious sources, which are merely correlated with the true sources but provide no additional information about the target when conditioning on these true sources.

The previous work [42] which compared multivariate TE and the greedy algorithm to the pairwise approach did so for standard time series of continuously varying signals sampled at a fixed interval. In this section, we verify that similar results hold when analysing event-based data using the continuous-time estimator of TE. Moreover, the analysis in this section will confirm the benefit of using the multivariate greedy approach, over the pairwise approach, when inferring networks from spike trains using TE.

We make use of the higher and lower synchrony simulations presented in Sec. II A, with 500 target spikes available. We applied a simple pairwise network-inference scheme to the resulting spike times from these simulations, which simply tested for statistically significant non-zero TE between each source-target pair. The resulting ROC curves are shown in Fig. 3. These ROC curves are created by sweeping through the *α* cutoff values (the threshold below which the *p* value must be for a link to be inferred) between 0 and 1 and recording the false-positive and true-positive rate observed at each *α* value. Calculating the *p* values for the statistical significance tests of non-zero TE was done by both counting the proportion of empirical surrogates (see Sec. IV B for a discussion of how these empirical surrogates are created) larger than the measured TE (Fig. 3f and Fig. 3b), as well as via a fitting a normal distribution to the surrogate values (Fig. 3h and Fig. 3d).

**FIG. 3:**
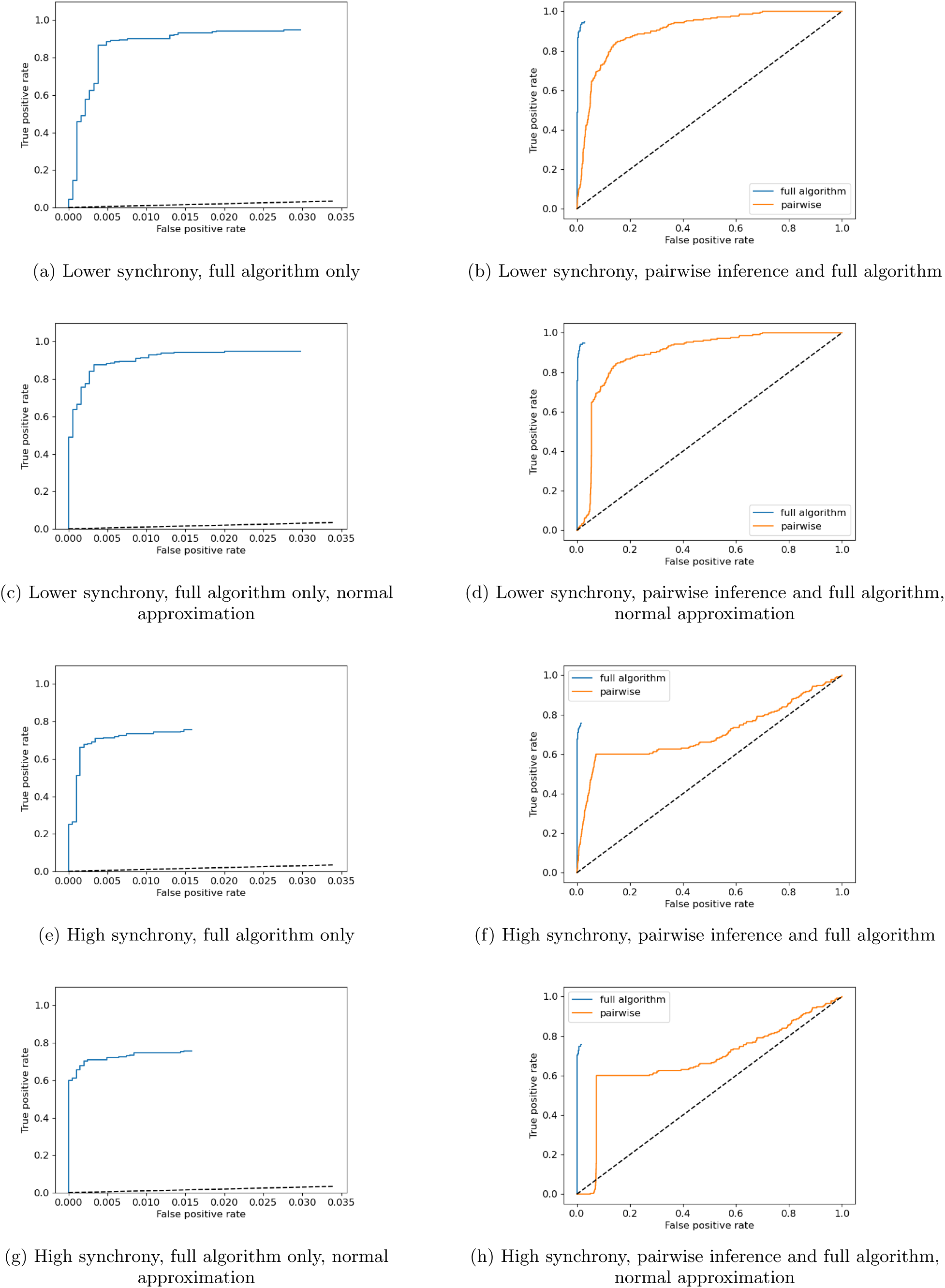
ROC curves for both the presented network inference technique, as well as performing the inference using the continuous-time estimator in a simple pairwise fashion. Plots are shown for the higher synchrony and lower synchrony examples from Fig. 1, with 1000 target spikes available to the algorithms. We also show plots for both the presented surrogate testing method as well as when using a normal approximation fitted to the surrogate population in order to estimate the *p* value.

The purpose of also performing this normal approximation is that it allowed us to use much lower *p* value thresholds for a given number of surrogate calculations than is possible when evaluating *p* values by counting proportions of empirical surrogates, making it possible to efficiently gain more resolution on the far left of the ROC curves.

We also ran the full greedy algorithm with different *α* cutoff values between 0 and 0.75, providing it with 1000 target spikes (the pairwise approach made use of the same number of target spikes). The final pruning step (see Sec. IV A) was, however, excluded, as this allowed for for greater computational efficiency in a single bulk run. The resulting ROC curves are also plotted in Fig. 3. Note that these ROC curves will not reach the point where the true positive rate and false positive rate both equal one, as we are unable to inspect *p* values larger than 0.75.

By inspecting the ROC curves in Fig. 3a through Fig. 3d we can compare the performance of the two approaches on the networks with lower levels of correlation. The full multivariate approach is seen to very quickly arrive at a true positive rate of above 0.9, for very few false positives, which again underlines the effectiveness of this approach. In contrast, the true positive rate of the pairwise approach rises much more slowly; in other words it costs a substantially larger number of false positives to achieve the same true positive rate. Visually we see this in that the ROC curve of the multivariate approach is markedly above that of the pairwise apprach.

We see an even starker difference in the performance of these two approaches when we look at the results of inference run on the networks exhibiting higher synchrony (Fig. 3e through Fig. 3h). Here, we see that the true positive rate of the pairwise approach rises much more slowly than the multivariate approach as before, but its performance also saturates around a true positive rate of around 0.6 before false positives begin to strongly dominate further inference. In this regime the entire population has a tendency to be active together and also remain quiescent together. As the activity of all neurons are therefore correlated, the pairwise approach is unable to delineate which particular neurons are driving the activity of others. The multivariate approach is, by contrast, more robust to the higher synchrony and still able to achieve very high true positive rates at incredibly low false positive rates.

These results demonstrate the substantial advantages in using multivariate TE estimation in conjunction with the greedy algorithm as opposed to a pairwise (functional network) approach.

### D. Inference of the effective networks of developing cell cultures

In order to demonstrate the utility of the application of this network inference scheme to biological data, we inferred the effective networks at various stages of development of cultures of dissociated cortical rat neurons. These recordings are part of a freely-available public dataset [31, 43]. See Sec. IV G for a summary of the nature of this dataset as well as details on how the network inference scheme was applied to it. In brief, cultures were allowed to develop over periods of around 30 days. On certain days, overnight recordings were performed. As these long overnight recordings contain sufficient numbers of spikes for effective application of information-theoretic estimators, they are eminently suitable for the application of our network inference approach. No spike sorting was performed, and so the networks are being inferred between the time series of the recording electrodes. This allows the nodes in the network to remain identifiable across different stages in development.

The results of applying the greedy algorithm along with the continuous-time estimator are displayed in Fig. 4. Note that the first recording days of each culture are not included in the figure as hardly any links (less than 10) were inferred in any of these recordings. We observe that effective networks with a rich and complex structure emerge, beginning to appear around the tenth day in vitro or so and quickly becoming more dense. This path of development correlates with the authors’ previous investigation [28] of these recordings, with simpler directed functional networks inferred using pairwise transfer entropy (via the same underlying estimator). The density of the networks inferred by the multivariate algorithm are lower than for the directed functional networks in [28], with the total number of edges in the last recording days declining from around 1000 to 2000 to around 100 to 200. This is common because the strongest action of the multivariate algorithm is to remove redundant sources [42].

**FIG. 4:**
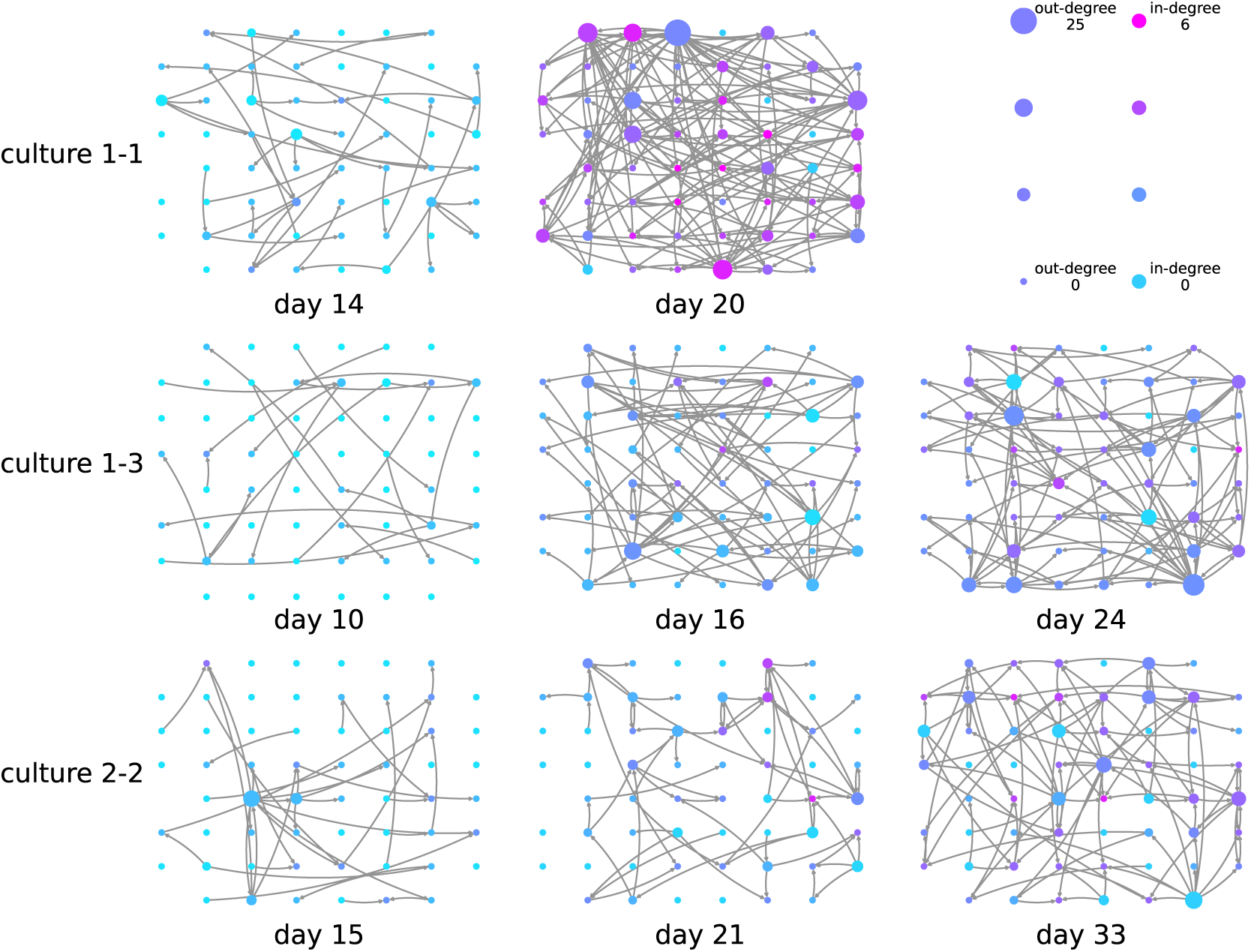
Effective networks inferred using the presented approach between electrodes in developing cultures of dissociated cortical rat neurons. Each node in the network visualisations is placed in the same relative spatial location that the corresponding electrode occupied in the recording apparatus. Networks were inferred at different stages of development (days *in vitro*). The recordings used are part of an openly-available public dataset [31, 43]. Node colour and size is proportional to the in and out degrees (see the legend in the top right). The spacing between the electrodes is 200 µm centre to centre [31, 43].

Despite the difference in density, the effective network structures retain some of the interesting features observed for the directed functional networks in [28], such as containing pronounced inward and outward hubs (that is, nodes with particularly high in-degree or out-degree), and various features of the networks being locked in early in development. Specifically, in the directed functional networks in [28] characteristics such as the total inward or outward information flow for a given node exhibited high correlation between early and late days of development. Here, Fig. 5 shows scatter plots of the out-degrees of the inferred effective networks on earlier and later days of development. We see in these plots that, as with the functional networks, in all cases, there is a positive correlation between the out-degree on the earlier and later days of development. Moreover, there are no statistically significant negative correlations. The positive correlation for the out-degree across the last two recording days is statistically significant for each culture, with some of these relationships, such as for culture 1-3, being particularly strong. Fisher’s method [44] for combining *p*-values was applied to all the *p*-values of the correlation coefficients (a *p*-value of 1 was used in the two cases of negative correlation). This results in a meta-test of the hypothesis that there is a positive correlation between the out-degree on the earlier and later days of development across the cultures. The null hypothesis of no or negative correlation was rejected at the 0.01 significance level.

**FIG. 5:**
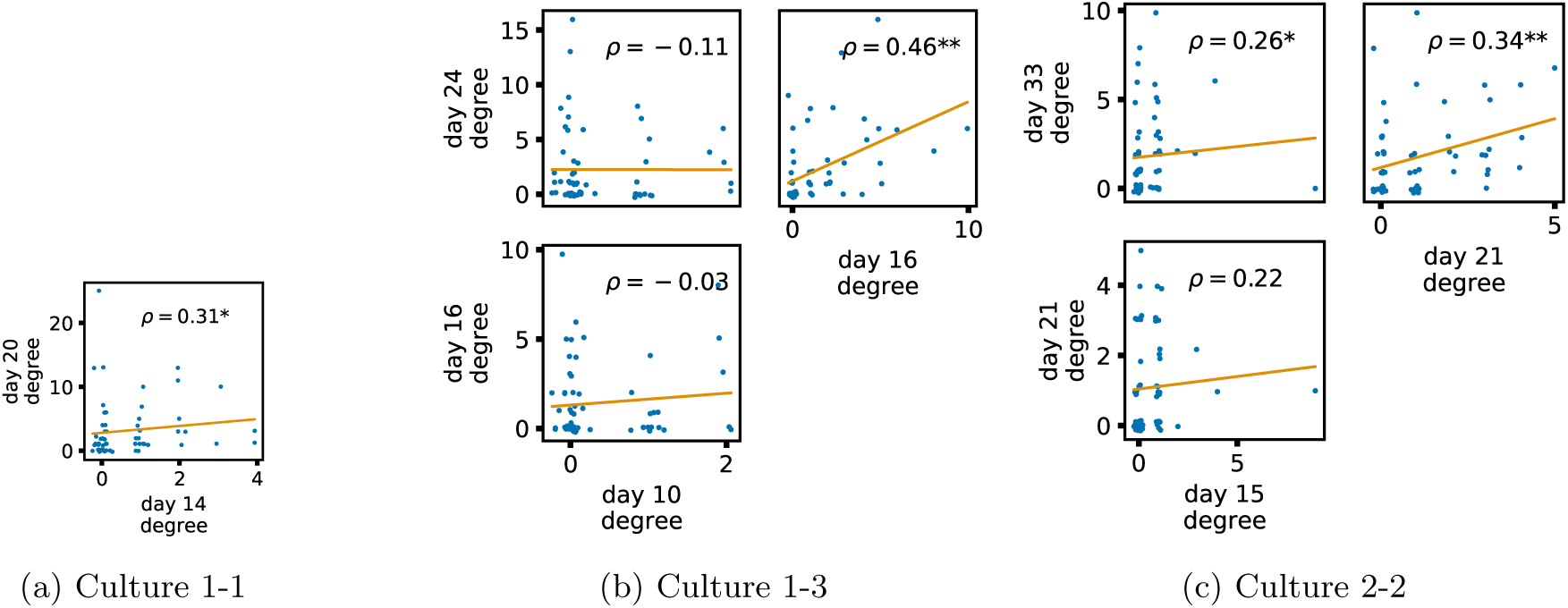
Plots showing the relationship between the out-degree of a given node over different days of development. Each group of plots shows scatter plots between all pairs of days for each culture analysed. Specifically, in each scatter plot, the *x* value of a given point is the out-degree of the associated node on an earlier day and the *y* value of that same point is the out-degree of the same node but on a later day. The days in question are shown on the bottom and sides of the grids of scatter plots. A small amount of Gaussian jitter (*σ* = 0.1) is added to the points to aid the visualisation of repeated values. The orange line shows the ordinary least squares regression. The Spearman correlation (*ρ*) between the out-degrees on the two days is displayed in each plot. Values of *ρ* significant at the 0.05 level are designated with an asterisk and those significant at the 0.01 level are designated with a double asterisk.

Fig. 6 shows similar plots, but for the in-degrees of the nodes. Again, we see a positive correlation between the in-degree on earlier and later days in every case. Fisher’s method for combining *p*-values was applied as above for the out-degree. The null hypothesis of no or negative correlation was rejected at the 0.01 significance level.

**FIG. 6:**
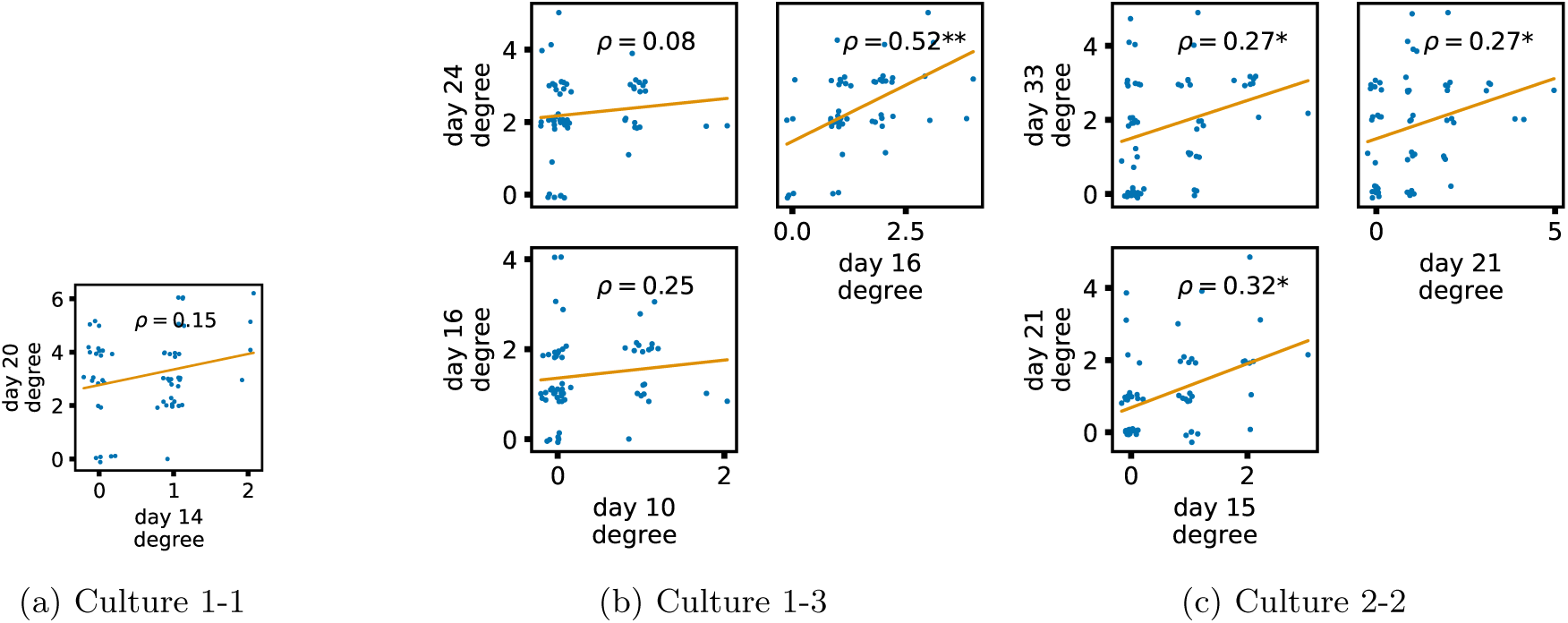
Plots showing the relationship between the in-degree of a given node over different days of development. Each group of plots shows scatter plots between all pairs of days for each culture analysed. Specifically, in each scatter plot, the *x* value of a given point is the in-degree of the associated node on an earlier day and the *y* value of that same point is the in-degree of the same node but on a later day. The days in question are shown on the bottom and sides of the grids of scatter plots. A small amount of Gaussian jitter (*σ* = 0.1) is added to the points to aid the visualisation of repeated values. The orange line shows the ordinary least squares regression. The Spearman correlation (*ρ*) between the in-degrees on the two days is displayed in each plot. Values of *ρ* significant at the 0.05 level are designated with an asterisk and those significant at the 0.01 level are designated with a double asterisk.

These results indicate that features of the effective networks, representing the multivariate information flows here, are being locked in early in development. This is particularly interesting since these are more sparse network models than the directed functional networks in [28], suggesting that the lock-in effect is deeply ingrained in the system.

Fig. 7 displays the proportion of possible links that are inferred in the networks at various physical distances between the nodes. Fig. 7a does so for the effective networks inferred in this work and Fig. 7b does so for the functional networks studied in the authors’ previous work. These plots show that the effective networks inferred on this dataset exhibit a clear preference towards links between nodes that are physically close together. This preference appears to become stronger with developmental time. By contrast, the functional networks do not exhibit this preference.

**FIG. 7:**
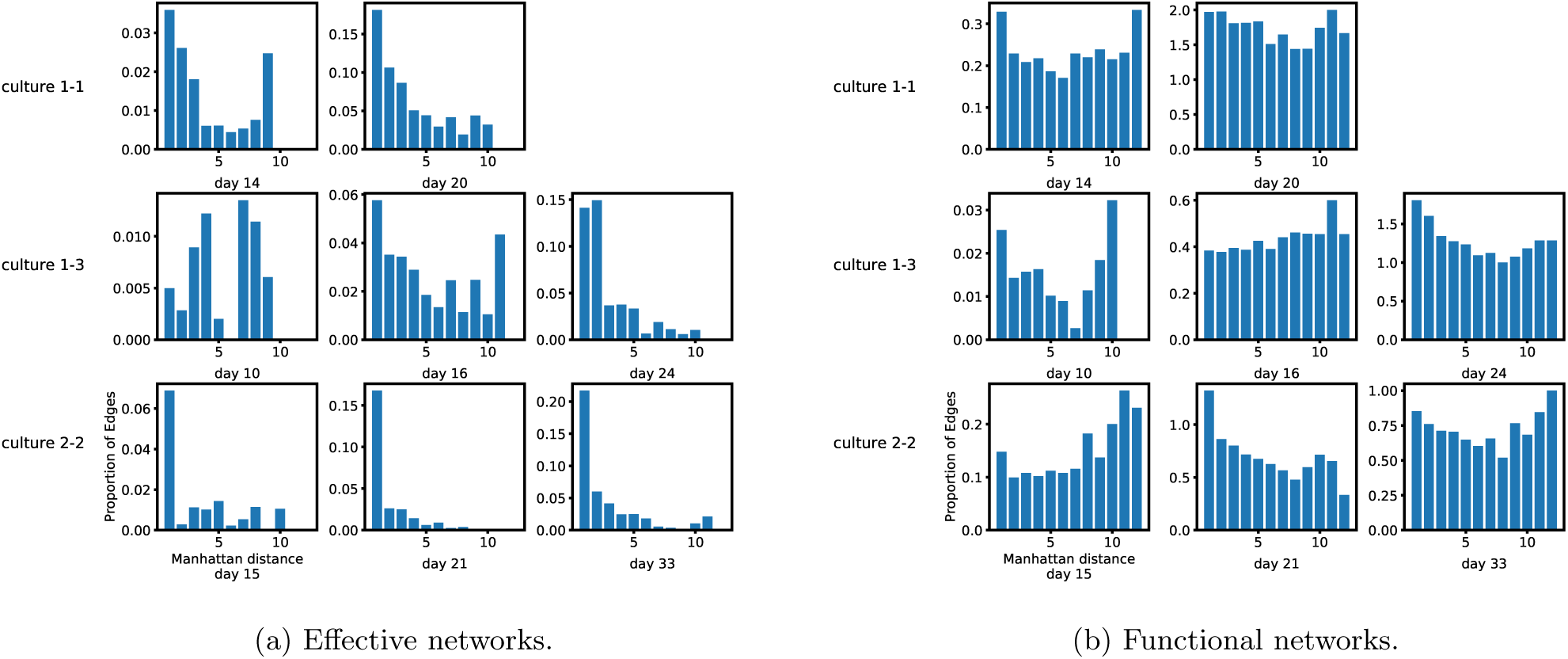
Bar plots showing the proportion of possible edges that were inferred at different inter-node distances. (a) shows the proportions for the networks inferred using the presented greedy algorithm and, whose diagrams are displayed in Fig. 4. (b) shows the same proportions for the functional networks inferred on the same data in the authors’ recent work [28]. The inference of such functional networks only considers pairwise relationships. The distances on the *x* axis are the Manhattan (cityblock) distances between electrodes. It is clear from the plots that, on this dataset, the effective network inference algorithm has a greater propensity to infer short distance links.

## III. DISCUSSION

In this work, we have validated the efficacy of the combination of an existing greedy multivariate TE-based network inference algorithm with a recently-introduced continuous-time estimator of TE on event-based data. Although we have validated the approach on neural spiking data, it can be applied to any event-based data, such as the times of social-media posts or the times of stock-market trades.

As the inference of networks from the spike times of neurons is a common goal within neuroscience, we expect this particular task to be a core application of the presented approach. Indeed, there have been several previous studies which have proposed TE-based methods for inferring networks from the spike times of neurons and evaluated them against ground truth [11–14]. There have also been a number of studies which used TE to infer networks from *in vitro* [15–21] and *in vivo* [22] recordings of spiking activity. These networks were found to exhibit highly non-random structure [16], including rich-club topologies [15]. All of this work has estimated the TE in a pairwise fashion, that is, without conditioning on other recorded processes. It has also performed estimation on discretised data. Networks inferred based on pairwise statistics are often referred to as *functional* (as opposed to *effective*) networks [5], since they are reporting pairwise relationships rather than a minimal multivariate directed model of the dynamics.

However, apart from a single very recent study [45], this previous work has always considered only the pairwise relationships between the activity on each node. As was demonstrated in Fig. 3, even when using the highly-effective continuous-time TE estimator, this approach suffers substantial drawbacks. Perhaps most notably, when the entire population is highly correlated, the pairwise approach is unable to find a small subset of sources which can collectively explain the activity of the target. Instead, it infers large numbers of sources due to the redundant information they hold with the true sources [42].

Here, by contrast, we have presented a multivariate approach to inferring *effective* networks from neural spike trains using TE. Unlike the pairwise approach, this strategy infers a set of parents for a target collectively rather than individually for each source. In doing so, the multivariate strategy considers the activity of other nodes within the network when determining the directed relationship between any two nodes. Specifically, as the multivariate strategy iteratively or greedily adds new candidate sources to the parent set for a target, it requires that each source provide statistically significant non-zero TE, when conditioning on all other current parents for the target. This is in contrast to the pairwise approach, which only requires a non-zero TE value between the source and target, without taking other processes into account. That iterative conditioning, along with a final associated pruning step, supports much more accurate inference because it eliminates redundant information from being spuriously attributed to other sources, and captures synergistic or collective interactions between multiple sources which jointly impact the target.

The term “accurate” here has specific meaning in the context in which we have evaluated the performance of the multivariate approach. Although we cannot and do not always expect the inferred effective networks to align with the underlying causal or structural network of the system being examined, under certain highly-specific idealised conditions (full observability etc., see Sec. I) we do indeed expect a minimal model explaining the dynamics of the variables (the effective network) to align with the causal structure in this way. As such, confirming such alignment – even though in highly-specific conditions – is an important validation of the performance of such an approach. Indeed, this validation is clear from our results, including in Fig. 1 and Fig. 2, and by the improved performance of the multivariate approach compared to pairwise, using the same underlying TE estimator, in Fig. 3. There, it was found that the multivariate approach was able to achieve a comparable true positive rate as the pairwise approach with a much lower corresponding false positive rate.

Maintaining a low false positive rate (or, equivalently, a high precision) is of utmost importance for network inference in a neuroscientific context. Zalesky states that “False positives are at least twice as detrimental as false negatives” [46]. As demonstrated in all of our results, the presented approach errs on the conservative side and consistently maintains high precision (a low false-positive rate) even in the challenging cases where the activity on the nodes is highly correlated (Fig. 1f). This high precision is a result not only of the multivariate strategy, but also the local permutation surrogate generation method used for the significance testing of the individual TE estimates being nonzero. This method was developed in tandem with the recently-developed continuous-time TE estimator [23] that is used here and was demonstrated to have substantially lower false-positive rates when compared with using traditionally-used approaches for surrogate generation in conjunction with this estimator.

Our results also compared the proposed approach with two existing approaches for the inference of connectivity from spiking data (CoNNECT in Fig. 8 and GLMs in Fig. 9), finding that it exhibited far superior precision in most cases, particularly so under conditions of high synchrony. The performance of the GLM approach degraded further when not using optimal parameters tuned using the ground truth (Fig. 10 and Fig. 11). This is particularly important when we reflect on the goal of effective network inference being to provide a “minimal model” that can explain the dynamics. Moreover, our approach, being grounded in information theory, has the unique advantage of inferring networks that are readily interpretable in terms of the fundamental computational operations of information storage, transfer and modification [7, 47]. And finally, as these measures can be estimated non-parametrically [48], they are not dependent on model assumptions and can capture any form of non-linear relationship.

**FIG. 8:**
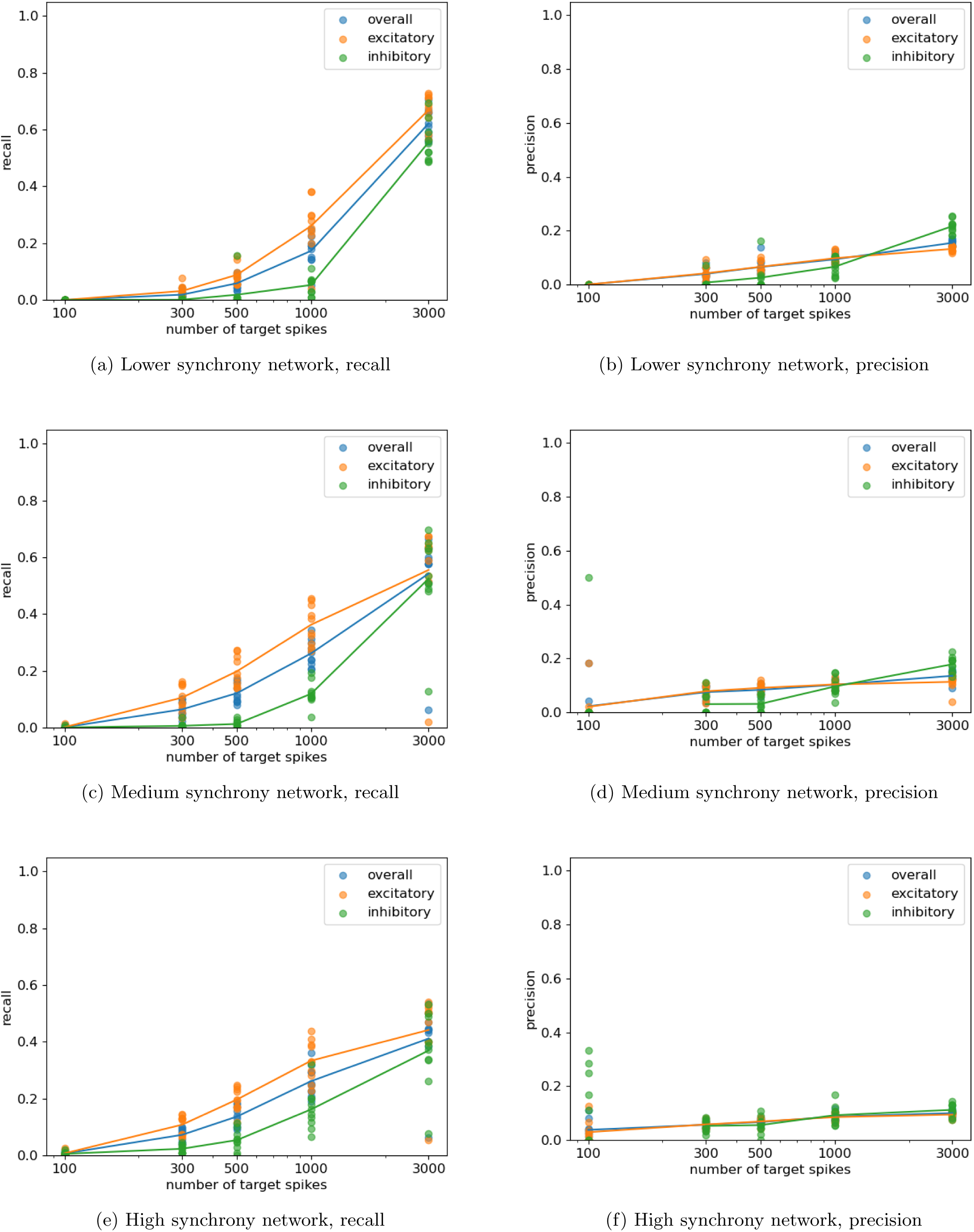
Plots showing the resulting precision and recall from running the CoNNECT algorithm [30] on networks of LIF neurons composed of 30 excitatory neurons and 20 inhibitory neurons. The ratio of the inhibitory to excitatory connection strength was varied in order to change the degree of synchrony in the network. Plots are shown for three different synchrony levels. Each plot contains points for the precision and recall for the inhibitory and excitatory neurons as well as the combined precision and recall. The lines pass through the means of these points.

Not only does this work present the first validation of TE for multivariate effective network inference on spike-train data, but it presents the first validation study of using the recently-developed continuous-time estimator of TE on event-based data such as spike trains in the context of network inference [23]. This estimator has been demonstrated to have many substantial advantages over the traditional discrete-time approach. These include consistency, lower bias and faster convergence. Of particular relevance to network inference, by representing the temporal pattern or history embeddings of processes using inter-event intervals, it is able to represent histories of substantial length using few dimensions and without any loss of time precision. This efficient use of dimensions facilitates building conditioning sets of significant size.

These various benefits culminate in a technique that is highly effective at the inference of networks from event times. We have demonstrated its strong performance, with high precision, in dynamical regimes ranging from low to high network synchrony (Fig. 1) as well as with varying levels of unobserved noise sources in the system (Fig. 2). Furthermore, high quality inferences were made with relatively low numbers of target spikes. In some instances (eg: Fig. 1a and Fig. 1b), near perfect reconstruction was achieved with only 3000 events per target.

These results all point to the strong potential for deploying this methodology in the inference of networks from recordings of the spike times of biological neurons. Tantalising hints of the results that might be expected were provided in Sec. II D. Further such applications remain a focus of future work. It is of particular interest to note that, in these effective networks, there was a strong preference towards inferring edges between nodes that are spatially close together, especially when compared with the functional network approach. This is likely due to the fact that effective networks are known to conform closer to the underlying structural networks than those inferred using pairwise, functional, methods [42]. This fits with the known fact that, in neural cell cultures, closer neurons are more likely to be connected [49]. However, despite this change in the topology of the networks, it is worth noting that the lock-in of information flows early in development, which was previously observed in functional networks [28], remained in these effective networks.

## IV. METHODS

### A. Greedy Algorithm

The greedy network inference algorithm used here was proposed in a range of papers [24–27], as summarised and studied in depth by Novelli et. al. [7], for traditional time series (a continuous-valued signal sampled at regular time intervals). We describe it here for completeness, and also to highlight some small changes that we made to adapt to the context of event-based data. The most significant change made is that only one inter-spike interval per source is considered as a candidate, and sequentially, from the most recent. This is as opposed to the original algorithm, where several lags from each source could be considered in the same selection round, and no ordering was imposed on the addition of these lagged samples.

The greedy algorithm is specified in Algorithm 1. We walk through its operation here, with reference to the line numbers in Algorithm 1.

We iterate over each process *R_i_* in the set of processes *R* (line 1). These processes are the raw timestamps of events (spikes). Each process is being considered as a target, for which the sources need to be inferred. It is worth noting that the computations performed for each target are considered completely independent of one another. As such, it is easy to parallelise this algorithm across the different targets. Indeed, such parallelisation was performed for the experiments presented in this paper.

We initialise a data structure to keep track of the last interval added to the conditioning set for each source and the target itself (line 2). This algorithm makes the assumption that more recent inter-event intervals from a given source (or the target itself) always have more influence over the target than inter-event intervals further in the past. Based on this assumption, inter-event intervals for a given source are only considered as candidates once more recent intervals for that source have been added to the conditioning set. As above, this is distinct from the operation of the algorithm for traditional time-series.

Before considering candidate sources, we determine the number of target history intervals to condition on. We always include at least one (the most recent) such interval. We continue incrementing the total number of intervals that we are conditioning on until the next interval does not provide a statistically significant reduction in the uncertainty of the target (line 4). This reduction in uncertainty is measured by the conditional Active Information Storage (AIS) [47], which is the mutual information between the last target history interval being considered and the current state of the target, conditioned on the more recent target history intervals. Note that this quantity can be easily estimated using the continuous-time TE estimator by simply considering this last target interval as a source interval. The active information storage is estimated on the original spike train and on *N*_surrogates_ surrogate processes (lines 5 and 6), constructed using the local permutation method described in Sec. IV E. The *p* value for the significance test is then the proportion of AIS estimates on the surrogates which are larger than the AIS estimate on the original process. If *p < p*_cutoff_, then the null hypothesis of zero AIS is rejected and the number of target intervals being added is incremented.

Returning to the canidate sources then, we continuously iterate parent selection for the target until the candidate source interval with the highest TE is no longer statistically significant (line 13). For each of these iterations, we iterate over every process other than the target under consideration (line 16). For each such candidate source, we estimate the TE between the most recent inter-event interval of that source that has not yet been added to the conditioning set and the target, conditioned on all intervals of all sources already added to the conditioning set (line 17). We then estimate the TE for *N*_surrogates_ surrogate processes between the same source and target and conditioned on the same conditioning set (line 18). We also bias-correct the original and surrogate TE estimates by subtracting the mean value of the surrogates from each estimate (lines 19 and 20).

Once this has been performed for every candidate source interval, we select the interval which had the highest bias-corrected conditional TE (line 22). We then estimate the *p* value associated with the null hypothesis of the conditional TE from this source interval being zero (line 23). This is done using the maximum statistic test (see Sec. IV B).

The above process continues until the selected candidate source interval (with maximum bias-corrected conditional TE) fails the significance test (line 13). The algorithm then moves onto the final pruning step, where it is checked that every source retains a statistically significant conditional TE, once conditioning on every other process added to the conditioning set. This step is necessary as a source added early in the process might be providing information about the target that is fully redundant with that held by sources added later in the greedy building of the conditioning set. Such redundant sources need to be removed.

To perform the pruning, we continually try removing source intervals from the conditioning set one-by-one, until every final source interval in the set is found to be statistically significant. In a mirror image to how the candidate intervals, for a given source, are added iteratively from the most recent and then further back in time, they are removed in order from the last interval to the most recent. In each round of pruning, we iterate over all sources which had an interval added to the conditioning set (line 31). We then estimate the conditional TE and associated surrogates for the last remaining added interval for that source (lines 32 and 33). We then calculate the *p* value corresponding to the null hypothesis (line 34) of zero TE in the normal manner (that is, not using the maximum statistic test). After iterating over all sources in the conditioning set, we then find the source index with the maximum *p* value (line 36). If this *p* value is greater than the specified *α* cutoff, then we remove the last added interval for that source from the conditioning set (line 39).

### B. Maximum Statistic Test

When considering adding sources to the conditioning set, we test the candidate source with the highest TE using the maximum statistic test (line 23 of Algorithm 1).

It is worth briefly describing the usual method for testing for non-zero TE using surrogates. We generate *N*_surrogates_ surrogates, which conform to the null hypothesis of no temporal relationship (zero TE), using a given surrogate generation algorithm (see Sec. IV E for a description of the surrogate generation method used here). We then estimate the TE on each of these generated surrogate series. The proportion of these estimates which are greater than or equal to the estimate on the original data is then an estimate of the probability that we would observe a value greater than or equal to what we estimated on the original data, under the null hypothesis of zero TE (and therefore it is our *p* value).

Novelli et. al. [7] highlighted the fact that using this test as is, when adding sources to the conditioning set, would lead to high false-positive rates. This is effectively a multiple comparisons problem, in that the test is being performed on the *maximum* estimated TE value from the set of candidate sources.

In order to compensate for this, we replicate the selection of the maximum candidate source in the significance testing step. Specifically, for each *i ∈ {*1, 2*, …, N*_surrogates_*}*, we compare the TE estimates on the *i*^th^ surrogate for each of the candidate sources. We select the maximum such estimate for each *i*. The population of surrogate values for the maximum statistic test is then made up of the resulting *N*_surrogates_ maximum values. The test then proceeds as normal.

#### Algorithm 1: Greedy TE algorithm for network inference from event-based data.

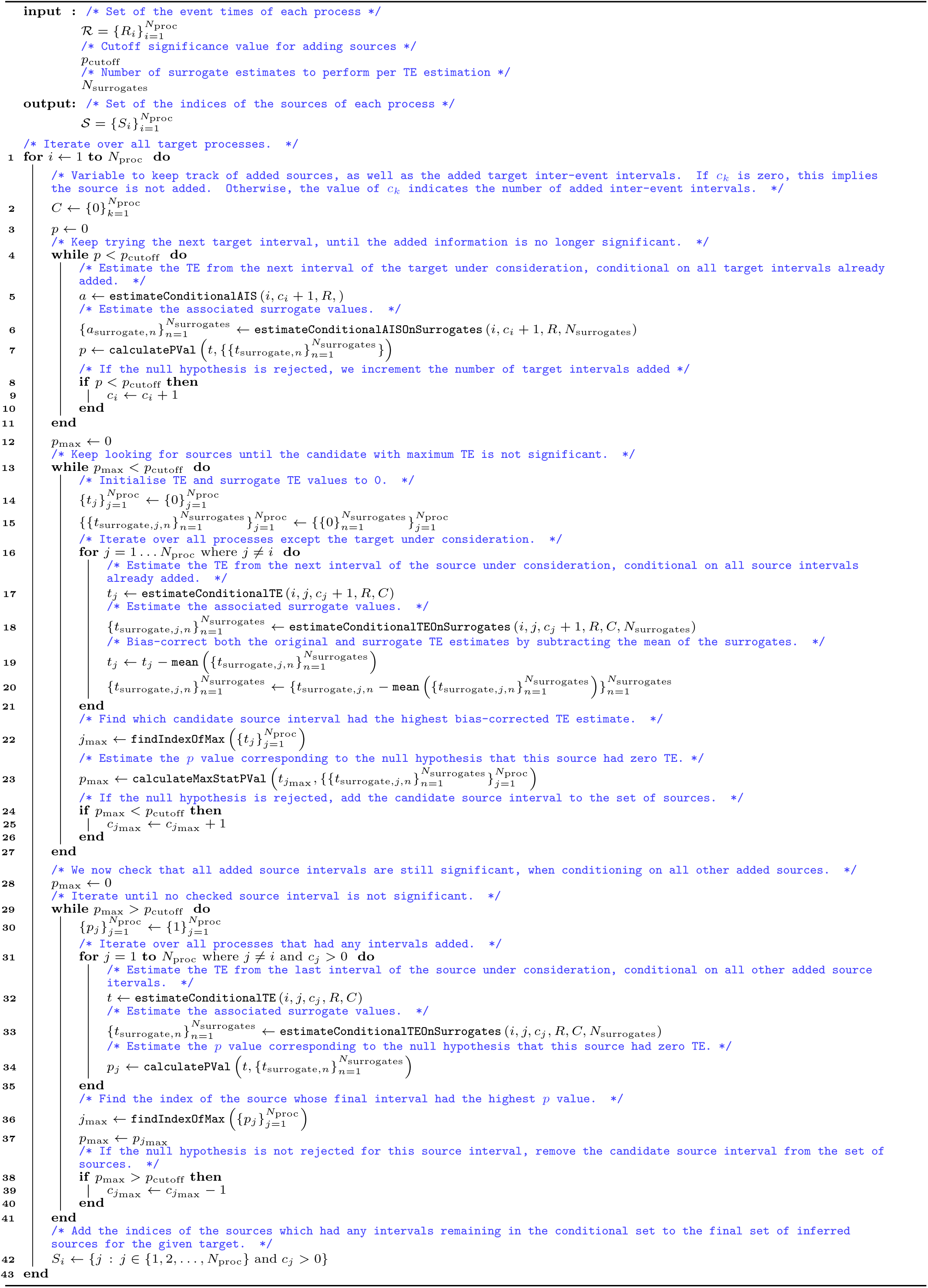

### C. Transfer Entropy Estimation – time-binning method

The transfer entropy [8, 9] is defined as the information provided by the past state or history embedding **y***_<t_* of a source variable (capturing its sequential temporal patterns) about the next value *x_t_* of a target variable, given the past state of that target **x***_<t_*. Specifically, this is a conditional mutual information:

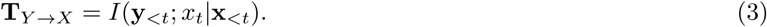

For standard time series processes *x_t_* and *y_t_*, the relevant past states are usually represented by Takens embedding vectors up to some embedding dimensions *l* and *k*, e.g. **y***_<t_* = *{y_t_*_−1_*, y_t_*_−2_*, …, y_t_*_−_*_l_}* and **x***_<t_* = *{x_t_*_−1_*, x_t_*_−2_*, …, x_t_*_−_*_k_}* (and potentially with embedding delays between the samples utilised as well); see [9]. The traditional method applying TE to spike trains operates by first discretising the spiking process into time bins (using some time bin width Δ*t*), providing a time series of binary values (spiking or not) on which the TE is then estimated. As above, the estimation of TE requires the use of embedding vectors of these binary time series to represent histories for the target, source and conditioning processes in the relevant conditional probability distributions for the target process. In order for these embeddings to both extend over a reasonable period of time and also capture fine subtleties in event timings, each embedding vector needs to consist of multiple time bins. Capturing effects occurring on both fine and large time scales is necessary as it is known that correlations in spike trains exist over distances of (at least) hundreds of milliseconds [50, 51]. Moreover, it is established that correlations at the millisecond and sub-millisecond scale play a role in neural function [52–55]. The use of multi-bin embedding vectors causes an explosion in the dimensionality of the state space over which probability distributions need to be estimated as conditioning processes are added, rendering the estimation of TE with substantial sets of conditioning processes infeasible. In the next section we discuss a recently developed TE estimator for spike trains which more efficiently handles the state space of interactions in such data.

### D. Transfer Entropy Estimation for spike trains in continuous time

It has, relatively recently, been shown that, for event-based data such as spike-trains, in the limit of small bin size Δ*t*, that the expected TE rate **T**^°^ *_Y_* _→*X*_ = lim_Δ*t*→0_ **^T^***^Y →X^* is given by the following expression [56]:

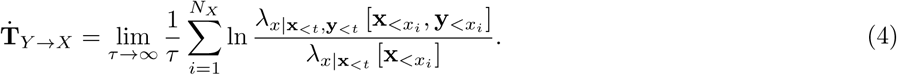

Here, *λ_x_*_|_**_x_***_<t,_***_y_***_<t_* [**x***_<xi_,* **y***_<xi_*] is the instantaneous firing rate of the target conditioned on the histories of the target **x***_<xi_* and source **y***_<xi_* at the time points *x_i_* of the spike events in the target process. *λ_x_*_|_**_x_***_<t_* [**x***_<xi_*] is the instantaneous firing rate of the target conditioned on its history alone, ignoring the history of the source. It is important to note that the sum is being taken over the *N_X_* spikes of the target during the sampling period *τ* : thereby evaluating log ratios of the expected spike rates of the target given source and target histories versus target histories alone, *when* the target does spike. As this expression allows us to ignore the “empty space” between events, it presented clear potential for allowing for more efficient estimation of TE on spike trains.

This potential was recently realised in a new continuous-time estimator of TE presented in [23], and all TE estimates in this paper were performed using this new estimator. In [23] it is demonstrated that this continuous-time estimator is far superior to the traditional discrete-time approach to TE estimation on spike trains. For a start, unlike the discrete-time estimator, it is consistent. That is, in the limit of infinite data, it will converge to the true value of the TE. It was also shown to have much preferable bias and convergence properties. Most significantly, perhaps, this new estimator utilises the inter-spike intervals to efficiently represent the history embeddings **x***_<xi_* and **y***_<xi_* in estimating the relevant conditional spike rates in (4).

This is in contrast with the traditional discrete-time estimator which uses the presence or absence of spikes in an array of time bins as its history embeddings (it sometimes also uses the number of spikes occurring in a bin). In order to avoid the dimensionality of the estimation problem becoming sufficiently large so as to render estimation infeasible, only a small number of bins can be used in these embeddings. To focus in on cell-culture data, previous applications of TE to this type of data have used a variety of bin sizes: 40 µs [15], 0.3 ms [11], and 1 ms [16, 19]. Some studies chose to examine the TE values produced by multiple different bin widths, specifically: 0.6 ms and 100 ms [17], 1.6 ms and 3.5 ms [20] and 10 different widths ranging from 1 ms to 750 ms [18]. And specifically, those studies demonstrated the unfortunate high sensitivity of the discrete-time TE estimator to the bin width parameter. Moreover, all of these studies have only used a single bin in the history embeddings. In the instances where narrow (*<* 5 ms) bins were used, only a very narrow slice of history is being considered in the estimation of the history-conditional spike rate. This is problematic, as it is known that correlations in spike trains exist over distances of (at least) hundreds of milliseconds [50, 51]. Conversely, in the instances where broad (*>* 5 ms) bins were used, relationships occurring on fine time scales will be completely missed. This is significant given that it is established that correlations at the millisecond and sub-millisecond scale play a role in neural function [52–55]. In other words, previous applications of transfer entropy to electrophysiological data from cell cultures either captured some correlations occurring with fine temporal precision or they captured relationships occurring over larger intervals, but never both simultaneously. This can be contrasted with the inter-spike interval history representation used by the continuous-time estimator. To take a concrete example, in the *in vitro* data we used, for the recording on day 24 of culture 1-3, the average interspike interval was 0.71 seconds. This implies that the history embeddings used are at least on average 0.71 seconds long, being longer than this in cases where multiple intervals are being used. This is despite the fact that our history representations retain the precision of the raw data (40 µs) and the ability to measure relationships on this scale where they are relevant (via the underlying nearest-neighbour estimators). Furthermore, the innovative representation of history embeddings as an array of inter-spike intervals allows for the application of the highly effective nearest-neighbour family of information-theoretic estimators [48, 57], which bring estimation efficiency and bias correction.

The challenges of using the discrete-time estimator only become more severe when one attempts to infer networks using conditional TEs. As there are now more processes being considered by the estimator (those in the conditioning set) the dimensionality of the estimation problem increases faster as we increase the embedding length. This places further pressure on keeping the number of bins in each embedding low, thus increasing the harshness in the tradeoff between history length and temporal precision. This is the likely reason behind the fact that almost all previous studies which evaluated the use of TE for the inference of spiking networks only made use of pairwise TE estimates [11–14]. This is as opposed to the multivariate conditional TE estimation used here, which takes into account the relationship of the target to other processes when considering its relationship to the given source.

The parameters used with this estimator for the simulated data are shown in Table I and those used for *in vitro* spike recordings are shown in Table II. The chief difference in the parameter values used in these situations is that, for the *in vitro* recordings a larger value of *k*_global_ (the number of nearest neighbours to consider in the initial searches) was employed (10 compared to 4). This was due to the observation that the estimates on the *in vitro* recordings exhibited much higher variance than those on the simulated data. It is a known property of nearest-neighbour information theoretic estimators that considering larger numbers of neighbours reduces their variance [48].

**TABLE I:** The parameter values used in the continuous-time TE estimator when used for the inference of the simulated spiking networks. A complete description of these parameters, along with analysis and discussion of their effects can be found in [23].

| Parameter | Description | Value |
| --- | --- | --- |
| $N_X$ | Number of spikes in the target spike train | varied (see text) |
| $k_{\text{global}}$ | Number of nearest neighbours to find in the initial search | 4 |
| $k_{\text{perm}}$ | Number of nearest neighbours to consider during surrogate generation | 20 |
| $N_U$ | Number of random samples of histories at non-spiking points in time | $20N_X$ |
| $N_{U,\text{surrogates}}$ | Number of random samples of histories at non-spiking points in time used for surrogate generation | $20N_X$ |
| $N_{\text{surrogates}}$ | Number of surrogates to generate for each node pair | 100 |

**TABLE II:** The parameter values used in the continuous-time TE estimator when used for network inference on the *in vitro* spike recordings. A complete description of these parameters, along with analysis and discussion of their effects can be found in [23].

| Parameter | Description | Value |
| --- | --- | --- |
| $N_X$ | Number of spikes in the target spike train | varied (see text) |
| $k_{\text{global}}$ | Number of nearest neighbours to find in the initial search | 10 |
| $k_{\text{perm}}$ | Number of nearest neighbours to consider during surrogate generation | 20 |
| $N_U$ | Number of random samples of histories at non-spiking points in time | $20N_X$ |
| $N_{U,\text{surrogates}}$ | Number of random samples of histories at non-spiking points in time used for surrogate generation | $20N_X$ |
| $N_{\text{surrogates}}$ | Number of surrogates to generate for each node pair | 100 |

As in the authors’ previous work applying the continuous-time TE estimator to *in vitro* spike recordings [28], a small change was made to the estimation procedure described in [23]. This was made in how random sample points were placed along the process both for the estimation of the TE and the generation of surrogates. Instead of laying out the *N_U_* and *N_U,_*_surrogates_ sample points randomly uniformly, we placed each one at an existing target spike, with the addition of uniform noise on the interval [*−*80 ms, 80 ms]. This was due to the fact that these recordings contain incredibly dense bursts. Such a sampling strategy is required in order to adequately sample these regions of intense activity.

An implementation of the estimator contained in the Java Information Dynamics Toolkit (JIDT) [58] software package was used in this study.

### E. Surrogate Generation

Surrogate processes were generated by applying an adaptation of the permutation method of Runge [59] to the spiking TE estimator, as detailed in [23]. In brief, this method permutes the history embedding vectors to destroy the relationship between the source intervals and the existence or absence of spiking in the target. However, it retains the relationship between the source history embedding intervals and the embedding intervals from the target and conditioning processes.

### F. Spiking Network Simulation

All network simulations were conducted using Leaky-Integrate-and-Fire (LIF) model neurons [34]. In this model, the membrane potential of the *i*^th^ evolves according to:

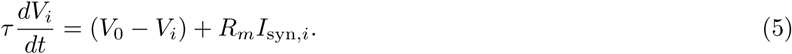

When *V_i_* crosses the threshold *V*_threshold_, the timestamp of crossing is recorded as a spike. *V_i_* is then set to *V*_reset_ and the evolution of the membrane potential is subsequently paused for the duration of the hard refractory period. *I*_syn*,i*_ is the synaptic input current into neuron *i*. Neurons were connected using alpha synapses [60]. Each synapse connecting neuron *j* to neuron *i* evolves according to:

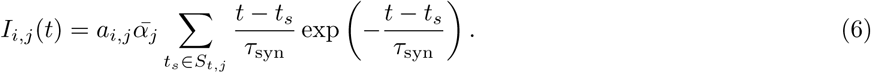

*A* is the connectivity matrix, with *a_i,j_* = 1 indicating that neuron *j* is a pre-synaptic input to neuron *i* and *a_i,j_* = 0 indicating otherwise. 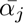 is the connection strength of the afferent connections from neuron *j*. All excitatory neurons share the same afferent connection strength 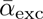. Inhibitory neurons, by contrast, have connection strength 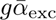. The sum is taken over the set of spike times in neuron *j* occurring before time *t*, *S_t,j_*. The synaptic current for neuron *i* is then the sum of the currents from all other neurons in the network, that is, *I*_syn*,i*_ = *I_i,j_*.

Each neuron was connected to exactly *n*_exc_ excitatory sources, chosen randomly from the set of excitatory neurons. Similarly, each neuron was connected to *n*_inh_ inhibitory sources, chosen randomly from the inhibitory sources. The specific parameter values used in the experiments described in Sec. II are shown in Table III and Table IV.

**TABLE III:** The parameter values used in the LIF network simulations at various levels of synchrony, presented in Sec. II A

| Parameter | Description | Value |
| --- | --- | --- |
| $N_{\text{exc}}$ | Number of excitatory neurons | 30 |
| $N_{\text{inh}}$ | Number of inhibitory neurons | 20 |
| $\tau$ | Membrane time constant | 20 ms |
| $R_m$ | Membrane resistance | 1 $\Omega$ |
| $V_{\text{reset}}$ | Membrane reset potential | 0 mV |
| $V_0$ | Membrane resting potential | 0 mV |
| $V_{\text{threshold}}$ | Spike threshold potential | 40 mV |
| $\tau_{\text{syn}}$ | Synaptic time constant | 20 ms |
| $g$ | Ratio of inhibitory to excitatory connection strength | $\{1, 1.5, 3\}$ |
| $\bar{\alpha}_{\text{exc}}$ | Excitatory connection strength | 20 mA |
| $n_{\text{exc}}$ | Number of excitatory sources per neuron | 3 |
| $n_{\text{inh}}$ | Number of inhibitory sources per neurons | 2 |
| $\lambda_{\text{stim}}$ | Rate of the Poisson stimuli | 200 Hz |
| $\bar{\alpha}_{\text{stim}}$ | Stimulus connection strength | 6 mA |

**TABLE IV:** The parameter values used in the LIF network simulations at various levels of synchrony, presented in Sec. II B

| Parameter | Description | Value |
| --- | --- | --- |
| $N_{\text{exc}}$ | Number of excitatory neurons | 30 |
| $N_{\text{inh}}$ | Number of inhibitory neurons | 20 |
| $\tau$ | Membrane time constant | 20 ms |
| $R_m$ | Membrane resistance | 1 $\Omega$ |
| $V_{\text{reset}}$ | Membrane reset potential | 0 mV |
| $V_0$ | Membrane resting potential | 0 mV |
| $V_{\text{threshold}}$ | Spike threshold potential | 40 mV |
| $\tau_{\text{syn}}$ | Synaptic time constant | 20 ms |
| $g$ | Ratio of inhibitory to excitatory connection strength | 3 |
| $\bar{\alpha}_{\text{exc}}$ | Excitatory connection strength | 17.5 mA |
| $n_{\text{exc}}$ | Number of excitatory sources per neuron | 3 |
| $n_{\text{inh}}$ | Number of inhibitory sources per neurons | 2 |
| $\bar{\alpha}_{\text{stim}}$ | Stimulus connection strength | 4 mA |

Each neuron also received an independent stimulus. In the experiments presented in Sec. II A, this source was an homogeneous Poisson point process. In the experiments presented in Sec. II B, it contained varying amounts of regularity. Specifically, in Sec. II B, each neuron received a stimulus with a spike rate *λ* drawn from a normal distribution with mean 500 hertz and standard deviation of 25 hertz. In the fully random case, the stimulus was generated as an homogeneous Poisson point process, with rate *λ*. In the fully regular case, spikes were placed at a fixed interval of 1*/λ*. In the semi-regular case, the spikes were placed with this same fixed interval, but gaussian noise with mean 0 and standard deviation 0.5 millisecond was added to each spike time. The connectivity strength between each stimulus and its target was specified by the parameter 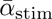.

### G. Analysis of *in vitro* Data

We made use of the same dataset as in the authors’ previous study [28] and analysed it in a very similar fashion. As such, the following section closely follows the discussion of this dataset in that previous work.

The spike train recordings used in this study were collected by Wagenaar et. al. [31] and are freely available online [43]. The details of the methodology used in these recordings can be found in the original publication [31]. A short summary of their methodology follows:

Dissociated cultures of rat cortical neurons had their activity recorded. This was achieved by plating 8×8 Multi-Electrode Arrays (MEAs), operating at a sampling frequency of 25 kHz with neurons obtained from the cortices of rat embryos. The spacing between the electrodes was 200 µm center-to-center. The MEAs did not have electrodes on their corners and one electrode was used as ground, resulting in recordings from 59 electrodes. In all recordings, electrodes with less than 100 spikes were removed from the analysis. This resulted in electrodes 37 and 43 being removed from every recording as no spikes were recorded on them. The spatial layout of the electrodes is available from the website associated with the dataset [43], allowing us to overlay the inferred networks onto this spatial layout as is done in Fig. 4.

Inspection of Figure 2 in the paper describing the original dataset [31] strongly suggests that these electrode recordings are sub-sampling the population of neurons. That is, spikes are only being detected from a fraction of the actual population. This is a possible cause of the low degree of the nodes in Fig. 4.

Recordings were conducted on most days, starting from 3-4 Days *In Vitro* (DIV). The end point of recording varied between 25 and 39 DIV. Longer overnight recordings were also conducted on some cultures at sparser intervals. In this work we make use of these longer overnight recordings. These recordings were split into multiple files. The specific files used, along with the names of the cultures and days of the recordings are listed in Table V. 30 Minute windows of spiking activity were extracted and used for network inference. Specifically, the number of target spikes *N_X_* was set as the number of spikes that fell within this 30 minute window for the given target neuron.

**TABLE V:** File numbers used for each culture on each day. These correspond to the file numbering used in the freely available dataset used in this study, provided by Wagenaar et. al.[31, 43]

|  |  |  |  |  |
| --- | --- | --- | --- | --- |
| **Culture 1-1** | day 4 | day 14 | day 20 |  |
|  | 2 | 2 | 2 |  |
| **Culture 1-3** | day 5 | day 10 | day 16 | day 24 |
|  | 2 | 2 | 2 | 2 |
| **Culture 2-2** | day 9 | day 15 | day 21 | day 33 |
|  | 2 | 2 | 2 | 2 |

The original study plated the electrodes with varying densities of cortical cells. However, overnight recordings were only performed on the ‘dense’ cultures, plated with a density of 2500 cells*/*µL.

The original study performed threshold-based spike detection by determining that a spike was present in the case of an upward or downward excursion beyond 4.5 times the estimated RMS noise of the recorded potential on a given electrode. The analysis presented in this paper makes use of these detected spike times. No spike sorting was performed and, as such, we are studying multi-unit activity (MUA) [61].

As the data was sampled at 25 kHz, uniform noise distributed between *−*20 µs and 20 µs was added to each spike time. This is to prevent the TE estimator from exploiting the fact that, in the raw data, inter-spike intervals are always an integer multiple of 40 µs.

### H. GLM Model

The implementation of Generalised Linear Models (GLMs) of spiking activity followed that of Song et. al. [40] very closely. We briefly list the few minor differences.

For the B-spline basis functions, we excluded all knot locations beyond 100 ms. This was done due to the membrane potential decay time constant (*τ*) in the simulated models being set to 20 ms (see Table IV), implying that statistical relationships beyond 100 ms would be very unlikely.

Song et. al. [40] propose finding the penalty weight parameter *λ* using the Bayesian information criterion (BIC), by iteratively trialling various penalty weight values. Performing this for each target spike train would have been computationally prohibitive given the large networks and long simulation times used in this work. Instead, this step was performed on a few trial runs and a single value of *λ* = 1*×*10^−3^ was chosen as it closely approximated that chosen by the BIC in all such trial runs.

We chose to designate the existence of a connection between a source and target when the GLM for the given target contained one or more non-zero weights assigned to the basis-splines associated with a given source.

Fitting of the GLM models was performed using the statsmodels [62] Python library.

## Appendix A: Comparison with CoNNECT Algorithm and Generalised Linear Models

Plots identical to those in Fig. 1, but showing the results of applying competing spiking network inference techniques to the same data. Specifically, Fig. 8 applies the CoNNECT algorithm [30], which makes use of pretrained convolutional neural networks to classify the existence (or otherwise) of edges between spike trains. Fig. 9 shows the results of applying a GLM model of spiking activity [40] to the data, and basing the inference of connectivity on the existence of non-zero weights in this model.

**FIG. 9:**
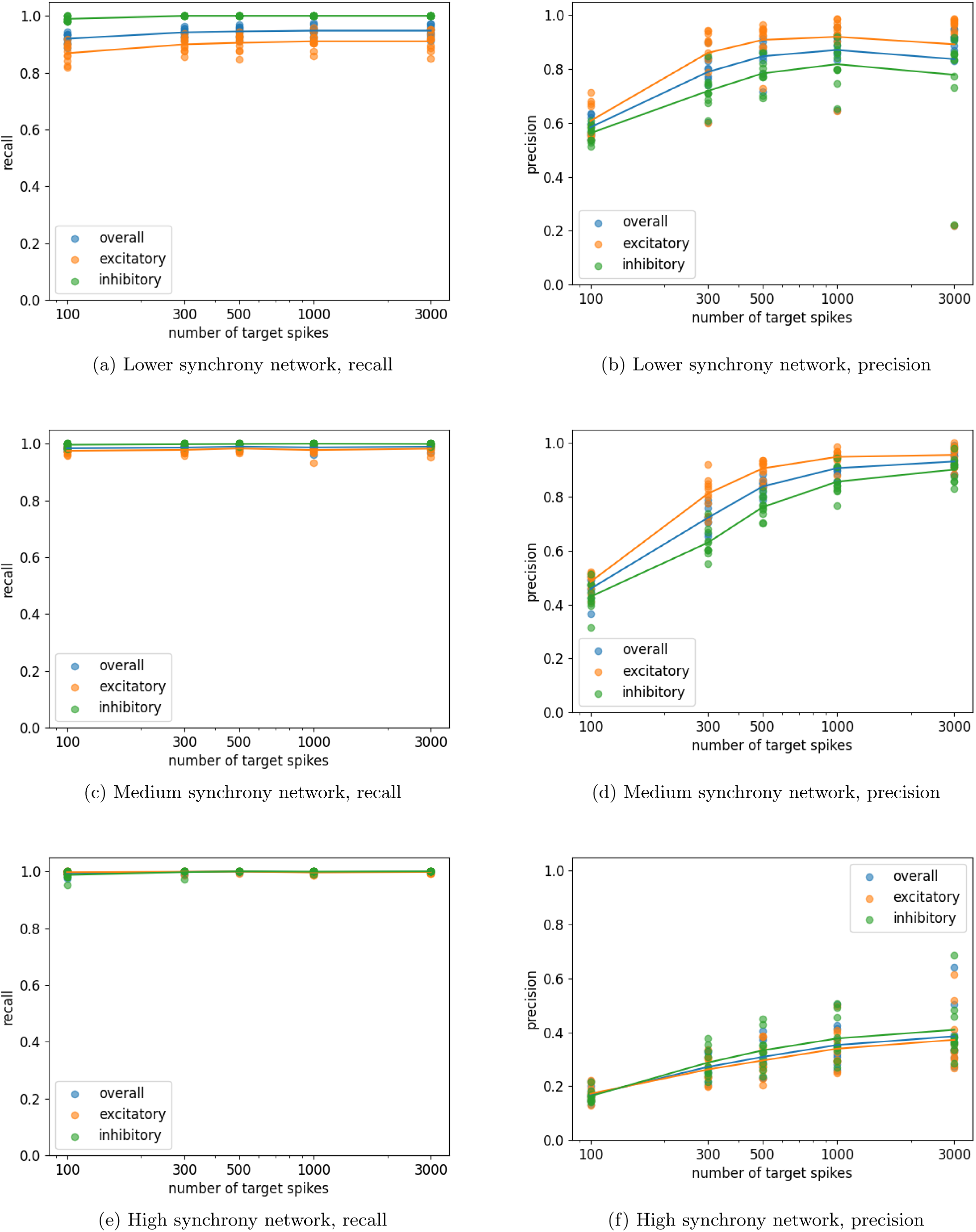
Plots showing the resulting precision and recall from using a GLM model of spiking activity [40] to infer the connectivity of networks of LIF neurons composed of 30 excitatory neurons and 20 inhibitory neurons. The ratio of the inhibitory to excitatory connection strength was varied in order to change the degree of synchrony in the network. Plots are shown for three different synchrony levels. Each plot contains points for the precision and recall for the inhibitory and excitatory neurons as well as the combined precision and recall. The lines pass through the means of these points.

**FIG. 10:**
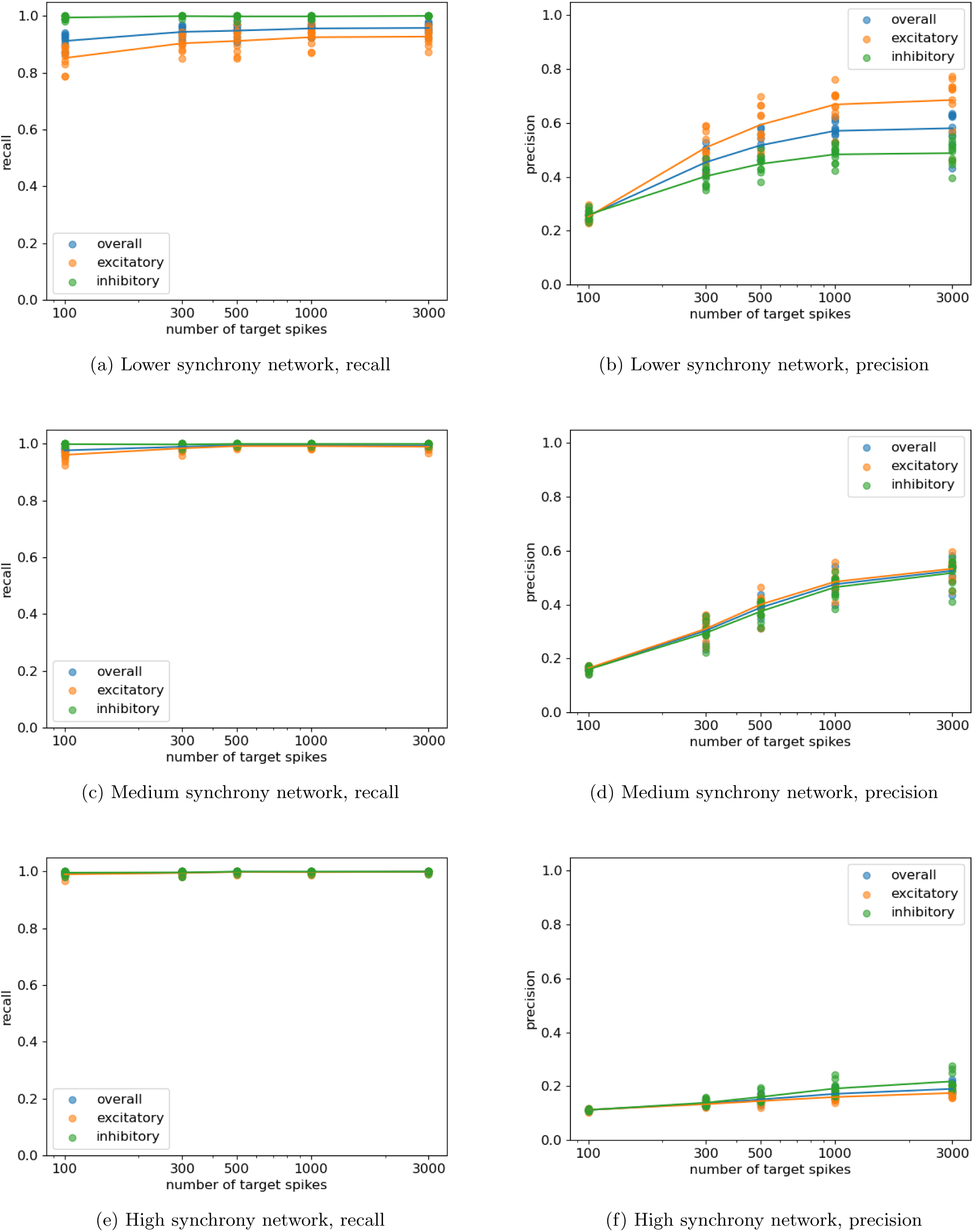
Identical plots to Fig. 9, but showing the use of a larger value of the time bin width Δ*t* (40 ms as oppose to 20 ms). There is a substantial drop in precision at this larger bin width. Note that, without access to the ground truth, there is no principled way to choose a best bin size. Although the results shown in Fig. 9 are quite promising, it is unlikely that they could ever be achieved in practice, as finding the optimal parameters (such as bin size) requires access to the ground truth.

**FIG. 11:**
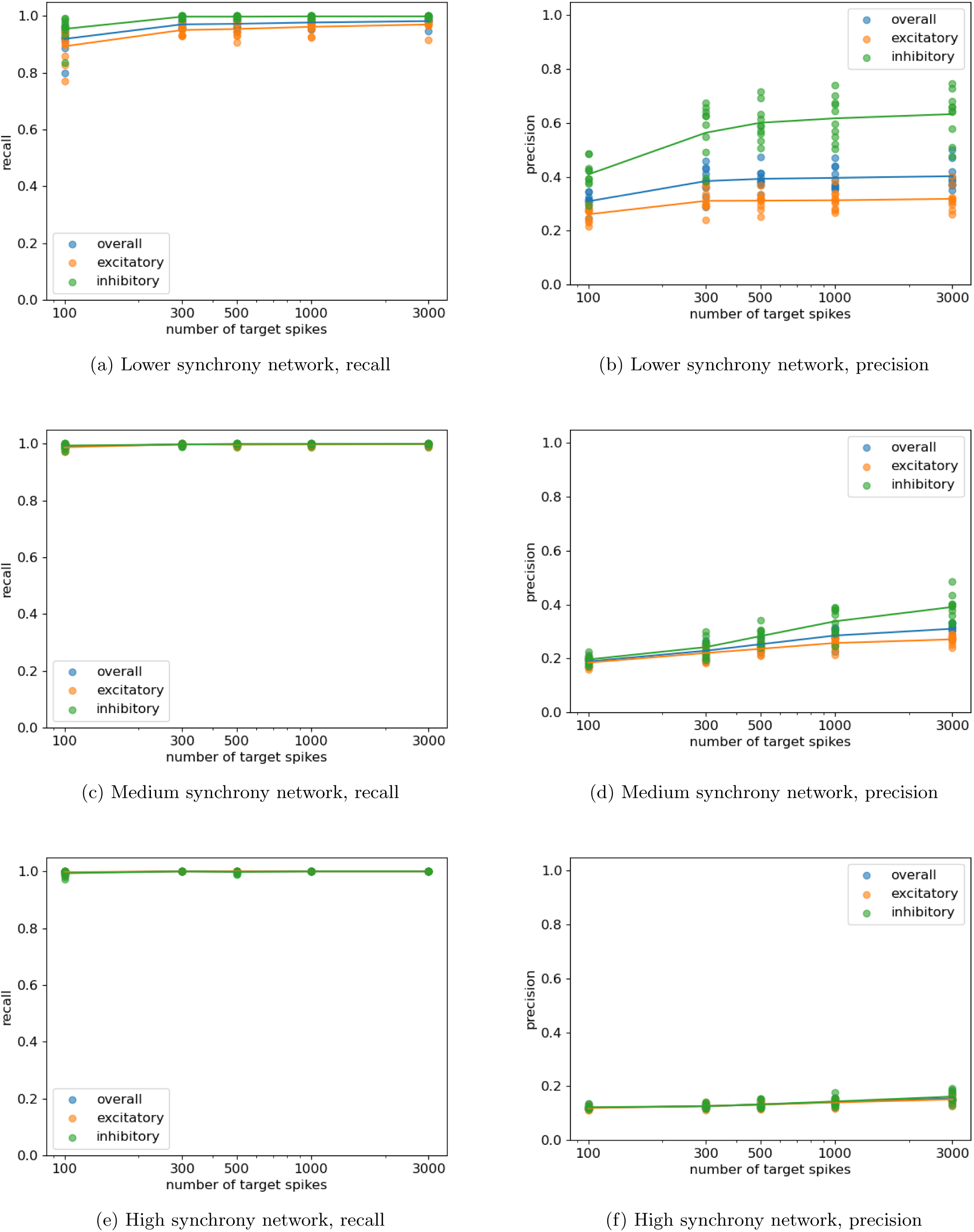
Identical plots to Fig. 9, but including B-spline knot locations beyond 100 ms. All 16 knot locations used Song et. al. [40] were incorporated, as opposed to only using the first 6, which was done elsewhere in this paper. Refer to Sec. IV H for more details on the use of B-splines in the GLM model. There is a substantial drop in precision when using these knot locations. Note that, without access to the ground truth, there is no principled way to choose the optimal knot locations. Although the results shown in Fig. 9 are quite promising, it is unlikely that they could ever be achieved in practice, as finding the optimal parameters (such as knot locations) requires access to the ground truth.

